# Two different mental disc forms in the Tashan cave *Garra:* different species or ecotypes?

**DOI:** 10.1101/2022.05.18.492420

**Authors:** Iraj Hashemzadeh Segherloo, Sajad Najafi Chaloshtory, Murtada D. Naser, Amaal G. Yasser, Seyedeh Narjes Tabatabaei, Gabriel Piette-Lauziere, Amirhossein Mashtizadeh, Amirmohammad Elmi, Omid Sedighi, Alieh Changizi, Eric Hallerman, Louis Bernatchez

## Abstract

In the Tashan Cave barb *Garra tashanensis*, inhabiting a small cave in south west Iran, two mental disc forms were observed. To assess their phylogenetic relationships, disc-less and disc-bearing individuals were analyzed using mitochondrial cytochrome oxidase subunit I (*COI*) partial DNA sequences. Both mental disc forms nested within one clade with absolute bootstrap support (*BS* = 100), and the genetic distances between the disc-bearing and disc-less individuals (0.3-0.7%) was considerably lower than inter-species mtDNA sequence distances reported among members of the genus *Garra*. Hence, the observed mental disc variation was not inferred to be a taxonomic feature or a consequence of character displacement. Instead, it was inferred as a case of character release to diversify among ecological niches in the limited subterranean habitat, which should be clarified in follow-up ecological and population genetic studies in more detail.

## Introduction

The mental disc – an extension of the lower lip in a number of labeonin fishes including *Garra* – has been used as a significant character for taxonomic and systematic inferences within the genus *Garra* (Hashemzadeh Segherloo et al. 2012, 2017; 2018; Mousavi-Sabet and Eagderi 2016). Based on mental disc characters, two phylogenetic groups had been considered for *Garra* spp., which include the *Garra variabilis* and *Garra rufa* clades (Hashemzadeh Segherloo et al. 2017); the members of *G. rufa* clade usually have a mental disc and the members of *G. variabilis* group lack the disc. In the clade with a mental disc (*G. ruffa*), the disc character has been used for description of the Lorestan cave barb *Garra lorestanensis*, which is sympatric with the blind Iran cave barb *Garra typhlops* in the Zagros Mountains of Iran (Hashemzadeh Segherloo et al. 2021; Musavi-Sabet and Eagderi 2016). This difference in mental disc of these species was also supported by mtDNA (Hashemzadeh Segherloo et al. 2012, 2017; Farashi et al. 2014) and genomic data (Hashemzadeh Segherloo et al. 2018). Furthermore, similar mental disc variation has been observed by the authors in the Tashan Cave barb *Garra tashanensis* population, which may be a taxonomically important character (character displacement) or alternatively, intra-species morphological diversification to exploit a broader ecological niche (character release).

Tashan Cave barb was described from a subterranean habitat (Tashan Cave, Behbahan, Khuzistan Province; N30°51’5” E50°10’35.7”) in karst formations of Southwest Iran (Mousavi-Sabet et al. 2016) in the Marun River drainage. This species is a member of the *G. rufa* clade which is also genetically highly diverged from extant congeners belonging the same group (*G. rufa* clade; see Musavi-Sabet et al. 2016 for more details). The species inhabits the Tashan Cave, which does not have any apparent connection to surface habitats. There are two morphotypes of the Tashan cave barb within the noted habitat: one with a well-developed mental disc and the other with no mental disc (Figure 1). Assuming the mental disc as a key feature of taxonomic significance, the individuals with no mental disc may be of different taxonomic status, similar to the case of the Iran and Lorestan cave barbs that originally were discriminated based on the mental disc character (Sargeran et al. 2008), otherwise, this may be a case of character release. In this study, both mental disc forms from the Tashan Cave are reported for the first time and compared using mitochondrial cytochrome C oxidase subunit I (*COI*) DNA sequences to infer their phylogenetic and taxonomic relationships.

**Figure 1.**
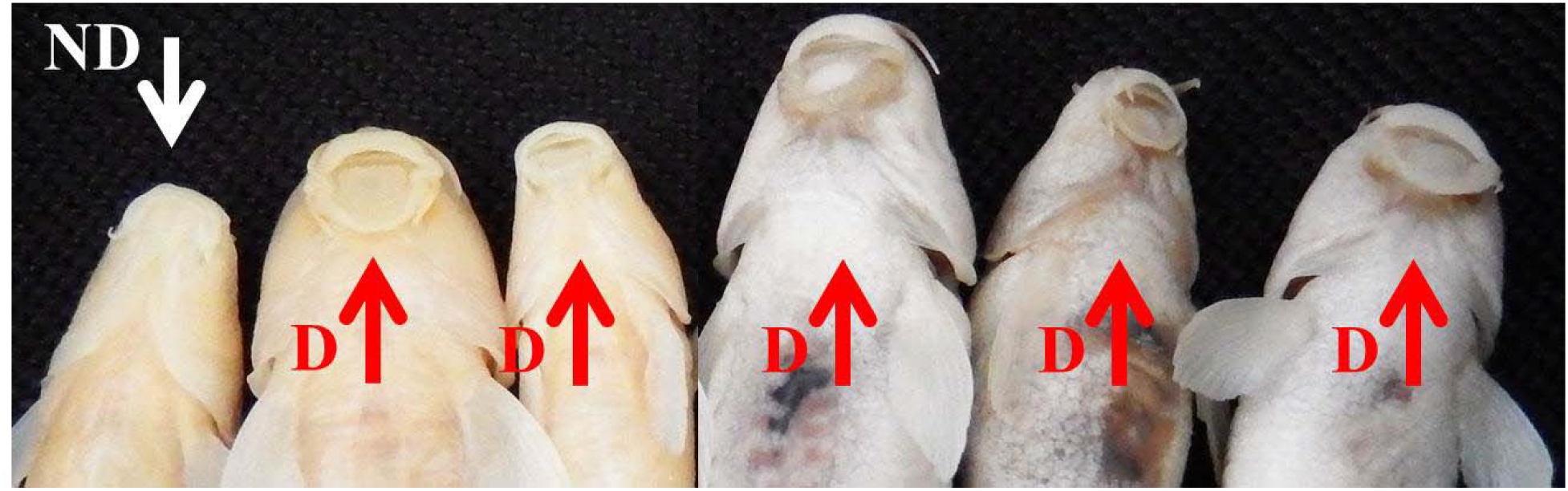
Ventral view of discless (white arrow, ND) and disc-bearing (red arrow, D) fish in Tashan Cave.

## Material and Methods

Due to conservation considerations, we collected ten individuals in February 2018 (*n* = 5) and September 2021 (*n* = 5) from the Tashan Cave using a dip-net (Figure 2). The specimens were preserved in 95% ethanol and after 24 hours, alcohol was refreshed. DNA was extracted using the salt extraction method of Aljanabi and Martinez (1997). Extracted DNA samples were quality-checked via electrophoresis through a 1% agarose gel.

**Figure 2.**
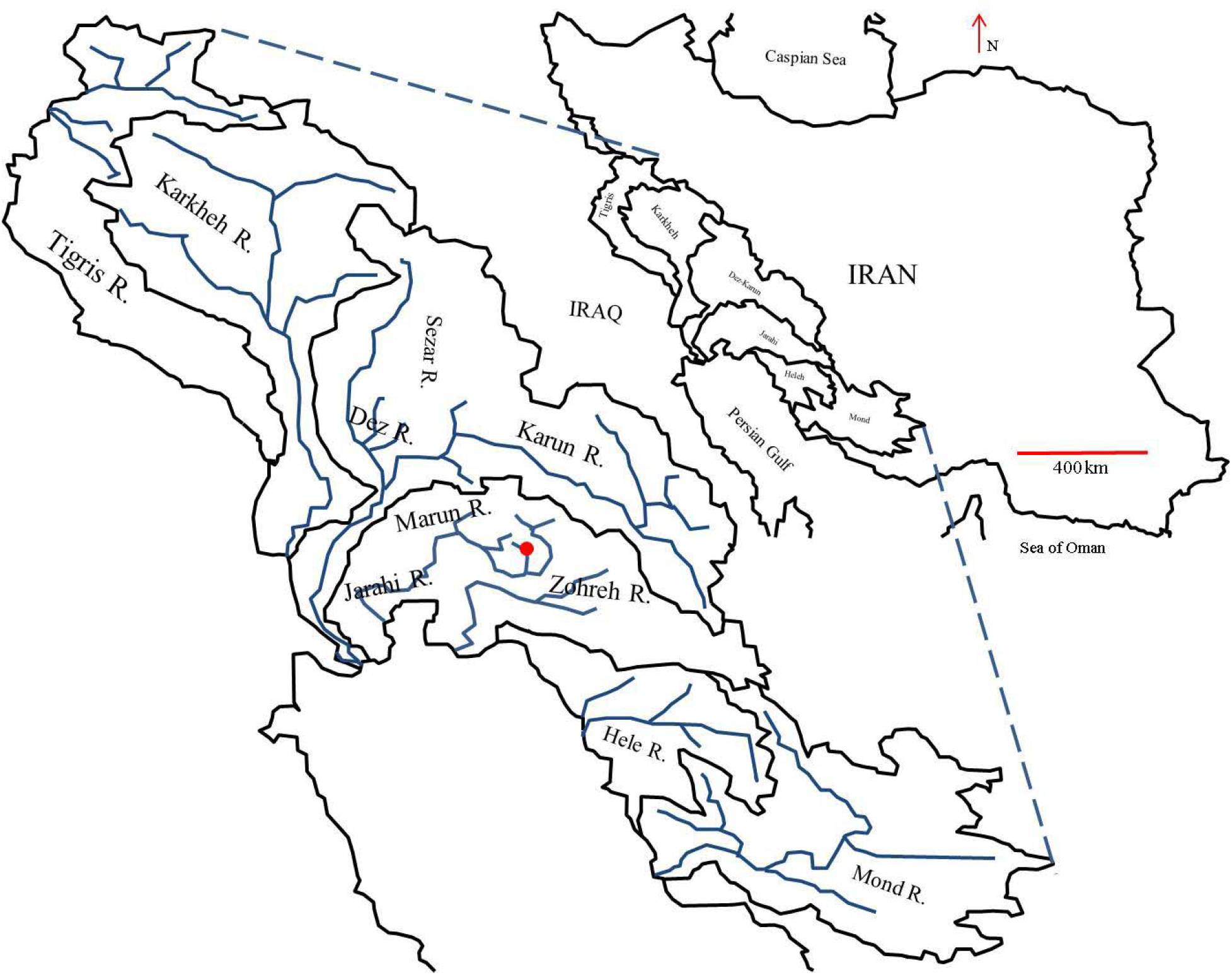
Map depicting the location of the Tashan Cave in southwest Iran. The red circle denotes the Tashan Cave.

The mitochondrial *COI* region was amplified using the *FishCOI-F:* 5’-AAYCAYAAAGAYATYGGYACCCT-3’ and *FishCOI-R:* 5’-TANACTTCNGGRTGNCCRAAGAAYCA-3’ primer set (Ivanova et al. 2007). Each 12.5-μl reaction included 6.25 μl Accustart II PCR mix (http://www.quantabio.com), 0.5 μl of each primer (10 μm), 3.25 μl deionized water, and 2 μl of DNA (10 ng/μl). The thermal profile for amplification included 60 s at 94 °C (preliminary denaturation); 30 cycles at 94 °C (30 s), 55 °C (30 s), and 72 °C (30 s); and one cycle at 72 °C (45 s). DNA sequencing was performed with the *FishCOI-F* primer using an ABI Prism 3130 sequencer (http://www.thermofisher.com) at the IBIS sequencing platform (Laval University, Quebec City, Canada; http://www.ibis.ulaval.ca).

The partial *COI* sequences were edited using BioEdit v. 7.2.5 (Hall 2014). The raw *COI* sequences were aligned using Muscle (Edgar 2004) with the default options in MEGA7 (Kumar et al. 2016). To find similar sequences for phylogenetic analyses, a BLAST (Johnson et al. 2008) search was performed in GenBank (NCBI). As some sequences from GenBank did not have the same lengths as our sequences, we used a common 532-bp sequence length for haplotype network and for Neighbour-Joining (NJ) and Maximum Likelihood (ML) phylogenetic tree reconstructions. For NJ phylogenetic tree reconstruction, the evolutionary distances were computed using the Kimura 2-parameter method (Kimura 1980), codon positions included were 1st+2nd+3^rd^, and all positions containing gaps and missing data (if any) were eliminated. To reconstruct the ML tree, the initial tree(s) for the heuristic search were obtained automatically by applying Neighbor-Join and BioNJ algorithms to a matrix of pairwise distances estimated using the Maximum Composite Likelihood (MCL) approach, and then selecting the topology with the superior log likelihood value. Codon positions included were 1st+2nd+3rd. All positions containing gaps and missing data were eliminated.

To provide support values for each node, 1,000 and 500 bootstrap replicates were run for NJ and ML tree reconstructions, respectively. To find the model that best described nucleotide substitution patterns, 24 different substitution models were tested using the model test implemented in MEGA7. The best model was selected according to the Bayesian information criterion (*BIC*). Other *Garra* spp. sequences from GenBank were used for reconstruction of the phylogenetic tree (Table 1). Three labeonin species – *Bangana ariza*, *Cirrhinus reba*, and *Labeo bata* – were used as out-groups. Kimura two-parameter (*K2P*) distances were calculated using MEGA7. DNA polymorphism indices were calculated using DNASP V 6.10.03 (Rozas et al. 2017).

**Table 1.** Mitochondrial *COI* sequences obtained from GenBank for use in phylogenetic reconstruction.

| Accession no. | Species | Accession no. | Species | Accession no. | Species |
| --- | --- | --- | --- | --- | --- |
| JX074145.1 | *Bangana ariza* | JF416298.1 | *Garra lorestanensis* | HQ235945.1 | *Garra tana* |
| MN862482.1 | *Cirrhinus reba* | KX983931.1 | *Garra orientalis* | KY365751.1 | *Garra tashanensis* |
| MG182184.1 | *Garra bispinosa* | MK074286.1 | *Garra ornate* | KY365751.1 | *Garra tashanensis* |
| JQ864601.1 | *Garra bourreti* | KM214807.1 | *Garra persica* | MN167175.1 | *Garra vinciguerrae* |
| MG182191.1 | *Garra cyrano* | MN255080.1 | *Garra rufa* | JX074212.1 | *Garra waterloti* |
| KF929909.1 | *Garra dembeensis* | KX983934.1 | *Garra surgifrons* | EU030664.1 | *Labeo bata* |
| MK572210.1 | *Garra lamta* | KX983934.1 | *Garra surgifrons* |  |  |

## Results

DNA of three out of the five specimens (one disc-less and two disc-bearing individuals) collected in 2018 were degraded, but the other seven individuals (three disc-less and four disc-bearing) had high quality DNA. The number of haplotypes (*nH*), haplotype diversity (*Hd*) and nucleotide diversity (*Pi*) calculated for the analyzed individuals (nine individuals: seven in this study and data of two specimens from Musavi-Sabet et al. (2016)) were: *nH* = 3, *Hd* = 0.417, and *Pi* = 0.003. The three haplotypes among the analysed specimens had a *K2P* distance of 0.57-0.76% (1-3 bp difference) from one another (Table 2, Figure 3). Two disc-less individuals and three disc-bearing individuals shared a haplotype with the two disc-bearing individuals reported by Mousavi-Sabet et al. (2016). The genetic distance of the analysed individuals from other congeners from Iran varied from 9.3-10.4% K2P distance.

**Figure 3.**
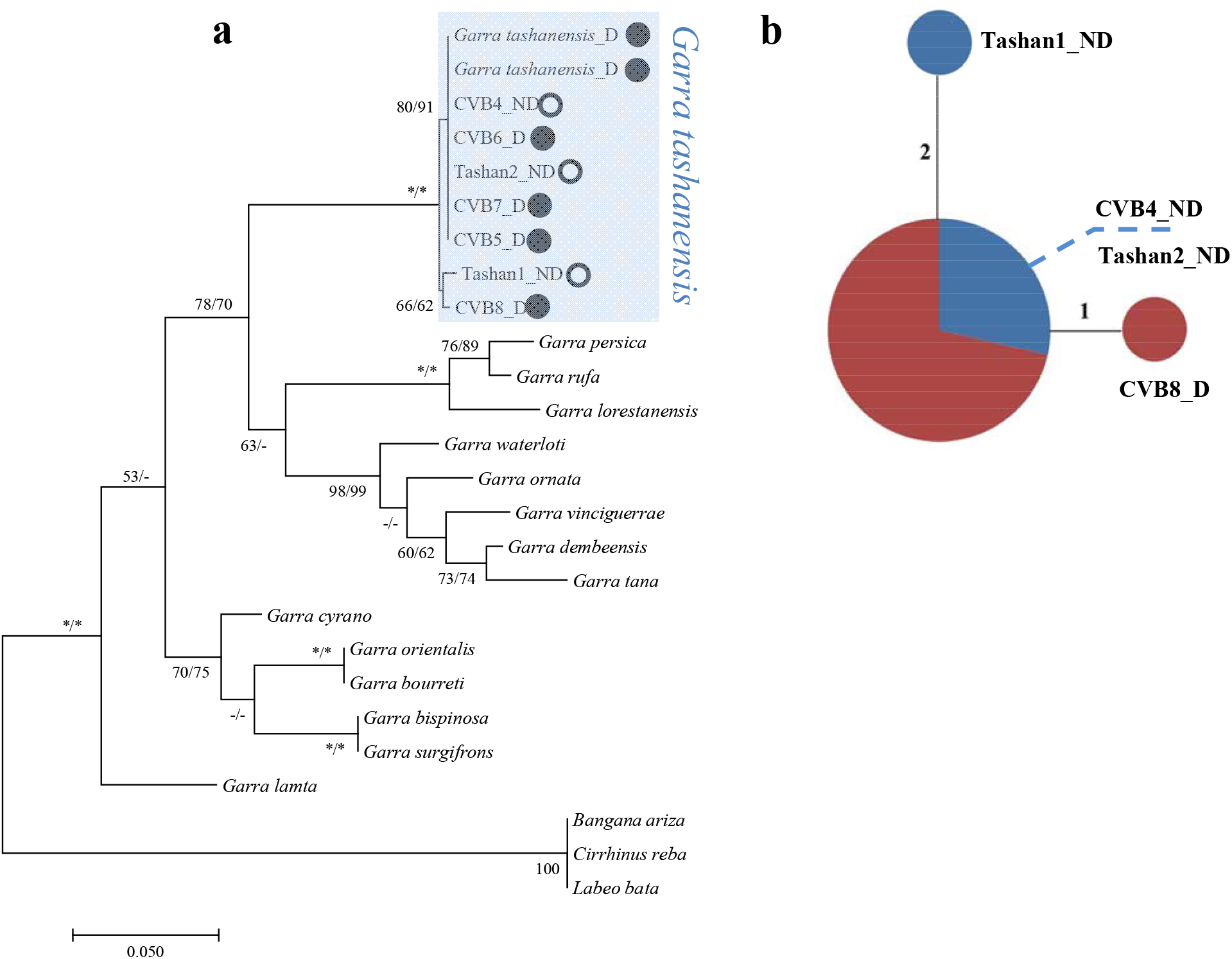
**a)** Maximum Likelihood (ML) phylogenetic three reconstructed for a 532-bp *COI* sequence for specimens collected for this study and sequences downloaded from GenBank. Bootstrap support for each branch are denoted beside branches: the values before the slash are the ML bootstrap values and after the slash are NJ bootstrap values. 100% bootstrap values are shown with an asterisk. Black-filled and white-filled circles respectively denote disc-bearing and disc-less *G. tashanensis*. b) Haplotype network of *G. tashanensis* sequences produced using POPART. Values along the lines between haplotypes indicate the number of mutational differences between each pair of haplotypes. The diameter of pie graphs is proportional to the frequency of haplotypes. Red color denotes disc-bearing individuals and blue color denotes disc-less individuals. The individual codes beside the pie graphs relate to codes of individuals in the phylogenetic network and identify disc-less individuals with their haplotypes and the disc-bearing individual with a haplotype that differs from the common haplotype.

**Table 2.** Estimates of evolutionary divergence between DNA sequences. Shown is the number of base substitutions per site between given sequences. Analyses were conducted using the Kimura 2-parameter model. Codon positions included were 1st+2nd+3rd. All positions containing gaps and missing data were eliminated. There was a total of 532 positions in the final dataset.

|  | Tash1_ND1 | TASH2_ND2 | CVB4_ND3 | CVB5_D | CVB6_D | CVB7_D | CVB8_D | *G. tashanensis_D* | *G. tashanensis_D* | *G. lorestanensis* | *G. rufa* | *G. persica* |
| --- | --- | --- | --- | --- | --- | --- | --- | --- | --- | --- | --- | --- |
| Tash1_ND1 |  |  |  |  |  |  |  |  |  |  |  |  |
| TASH2_ND2 | 0.76 |  |  |  |  |  |  |  |  |  |  |  |
| CVB4_ND3 | 0.76 | 0.00 |  |  |  |  |  |  |  |  |  |  |
| CVB5_D | 0.76 | 0.00 | 0.00 |  |  |  |  |  |  |  |  |  |
| CVB6_D | 0.76 | 0.00 | 0.00 | 0.00 |  |  |  |  |  |  |  |  |
| CVB7_D | 0.76 | 0.00 | 0.00 | 0.00 | 0.00 |  |  |  |  |  |  |  |
| CVB8_D | 0.57 | 0.57 | 0.57 | 0.57 | 0.57 | 0.57 |  |  |  |  |  |  |
| *G. tashanensis_D* | 0.76 | 0.00 | 0.00 | 0.00 | 0.00 | 0.00 | 0.57 |  |  |  |  |  |
| *G. tashanensis_D* | 0.76 | 0.00 | 0.00 | 0.00 | 0.00 | 0.00 | 0.57 | 0.00 |  |  |  |  |
| *G. lorestanensis* | 10.48 | 10.03 | 10.03 | 10.03 | 10.03 | 10.03 | 10.24 | 10.03 | 10.03 |  |  |  |
| *G. rufa* | 9.80 | 9.36 | 9.36 | 9.36 | 9.36 | 9.36 | 9.56 | 9.36 | 9.36 | 4.11 |  |  |
| *G. persica* | 9.36 | 9.36 | 9.36 | 9.36 | 9.36 | 9.36 | 9.56 | 9.36 | 9.36 | 4.52 | 1.92 |  |

The optimal NJ phylogenetic tree had a total branch length of 0.472. Based on the Bayesian Information Criterion (*BIC*), the model that best described the nucleotide substitution pattern for our data-set was the Hasegawa-Kishino-Yano (HKY) model with a discrete Gamma distribution (5 categories (+*G*, parameter = 0.1688); see Supporting Data). Among the trees reconstructed, we used the tree with the highest log likelihood (−1955.4988). Both the NJ and ML trees were similar in topology; we present the ML tree with bootstrap support values drawn from both phylogenetic reconstruction approaches (Figure 3). On both the NJ and ML phylogenetic trees, all the analysed individuals nested in a shared phylogenetic group with *G. tashanensis* with absolute bootstrap support (BS = 100; Figure 3). No discrete clustering pattern was observed between the disc-bearing and disc-less individuals.

## Discussion

### The origin of Tashan Cave barb

As noted above, Tashan Cave is located in the Marun River drainage, which is adjacent to the Zohreh River drainage. Hence, the most probable source for colonization of Tashan Cave is the Marun River or its tributaries. As reported in Hashemzadeh Segherloo et al. (2017) and Kiani et al. (2017), the counterparts of *G. tashanensis* in the Marun and Zohreh rivers are *G. rufa*. The high sequence distance between *G. tashanensis* and *G. rufa* or other *Garra* spp. (*K2P* distance > 9%) in neighbouring river drainages refutes any possible recent colonization of the cave by the modern surface-dwelling *Garra* spp. of the region. It is noteworthy that the modern *Garra* spp. of the region have been affected by invasion of *G. rufa* during the last glaciation when river drainages in the Persian Gulf Basin were connected to the Tigris River (Fagan 2014). For example, *G. mondica* in the Mond River drainage, *G. gymnothorax* in the Karun, Dez, and Karkheh river drainages, and *G. rufa* in the Tigris River drainage that are the modern inhabitants of the surface waters in the Persian Gulf Basin, all show high mtDNA sequence divergence along with high genomic (nDNA) similarity (Hashemzadeh Segherloo et al. 2018), which is probably an indication of the noted colonization event by *G. rufa*. No close relatives of *G. tashanensis* has been reported in surface waters of the Persian Gulf basin (Hashemzadeh Segherloo et al. 2017, 2018; Kiani et al. 2017). The case of *G. tashanensis* is similar to that of *G. lorestanensis* and *G. typhlops* or *Eidinemacheilus smithi* and *Eidinemacheilus proudlovei* (Feyhof et al. 2016; Hashemzadeh Segherloo et al. 2016, 2017) for which no closely related surface-dwelling congeners have been reported. This current situation probably is the consequence of ancient extinctions of the ancestral populations via drought periods and later colonization events by species including *G. rufa* and *G. gymnothorax*. In such cases, the subterranean habitats may have proven stable refuges for *Garra* spp. to survive environmental extremes and probably ecological invasions due to their stable environmental conditions (Davis et al. 2013).

### Signatures of limited population size

The haplotype diversity in *G. tashanensis* is lower than those reported for *G. lorestanensis* and *G. typhlops* (Hashemzadeh Segherloo et al. 2012; Farashi et al. 2014). Farashi et al. (2014) reported 11 *Cyt-b* haplotypes among 16 *G. lorestanensis* and *G. typhlops* they analysed, and Hashemzadeh Segherloo et al. (2012) reported three *COI* haplotypes for six specimens they studied. In comparison, for nine *G. tashanensis* analyzed in this study and in Mousavi-Sabet et al. (2016), only three *COI* haplotypes are reported. Lower haplotype diversity of *G. tashanensis than for G. typhlops* and *G. lorestanensis* may be due to its smaller population size. Bagheri et al. (2016) reported a population size of 300-600 individuals for *G. typhlops* and *G. lorestanensis*, and Mousavi-Sabet et al. (2016) reported a population size of at least 60-100 individuals in the two pools of 20-30 m^2^ surface area in Tashan Cave. Compared to the nuclear genome (nDNA), the mitochondrial genome (mtDNA) has an effective size 25% that of the nDNA effective size (Hallerman 2003); this lower effective size makes mtDNA diversity erode more rapidly in response to random genetic drift or population bottlenecks, which can be an explanation for low mitochondrial haplotypic diversity of *G. tashanensis* in Tashan Cave.

### Character displacement or release?

The existence of two sympatric *Garra* species with differing disc morphologies has been interpreted as a possible case of character displacement (Hashemzadeh Segherloo et al. 2017; 2021), which allows the respective species to use different ecological niches and reduce inter-species competition in sympatry (Schluter 2000; Gray et al. 2005; Pfennig and Pfennig 2010; Robinson and Pfennig 2013). However, our results do not support the existence of phylogenetically distinct *Garra* species in Tashan Cave, since there is no important mtDNA differentiation associated with mental disc variation and most likely with speciation (see Zamani-Faradonbeh et al. 2021 for more details on inter-species sequence distances among Iranian *Garra* spp.). At the intraspecific level, when no other closely related species is present in the habitat, it is possible that a number of individuals from the same population undergo morphological changes in their characters to diversify niches available to the species, i.e., character release (Robinson and Wilson 1994). According to the data presented in previous studies on other Iranian subterranean fish, it appears that mental disc morphology may not always be considered a stable taxonomic character. For example, Hashemzadeh Segherloo et al. (2013) reported an Iran blind cave barb (*G. typhlops*) with a mental disc, while the species had been known to have no mental disc. Our unpublished genomic data show that the disc-bearing *G. typhlops* reported in Hashemzadeh Segherloo et al. (2013) was not a hybrid individual with *G. lorestanensis*, the disc-bearing cave barb sympatric with *G. typhlops*. This again indicates variation of the mental disc character within the genus *Garra. Garra tashanensis* inhabits a limited subterranean habitat of around 50 m^2^ (Musavi-Sabet et al. 2016) in a semi-arid region where groundwater resources are limited (Ashraf et al. 2021). The Tashan Cave habitat is limited to two small, stagnant pools located 500 m from each other during summer (Mousavi-Sabet et al. 2016). Musavi-Sabet et al. (2016) did not report any flowing aquatic habitat in the Tashan Cave. Assuming such habitat limitations and other limiting factors that exist in cave habitats, it is possible that the mental disc variation in *G. tashanensis* to be an evolutionary response based on standing genetic variation to the limited energy resources and limited habitat area, expression of which increases the likelihood of sustaining a larger population size with lower levels of intraspecific competition i.e., character release.

### Distribution pattern

Hashemzadeh Segherloo et al. (2018) reported that *G. lorestanensis* – a mental disc-bearing cave barb species – only appears during the fluvial periods of the year when water flows in the habitat, and *G. typhlops* – the disc-less species - is observed during most of the year independent of water flow regime. The samples in this study were collected in summer and late winter i.e., in dry and fluvial periods of the year, but no differential frequencies of disc-bearing and disc-less individuals was observed in Tashan Cave; in both sampling periods, we collected both disc-bearing and disc-less individuals. Clarification of factors driving expression of mental disc-bearing and discless morphs in Tashan Cave awaits more-detailed ecological and population genetic studies.

## Acknowledgments

This work is dedicated to Mesdames Madeleine Drouin and BöyükKhanim Ahmadi Segherloo. This work was supported by a NSERC (Canada) Discovery grant (http://www.nserc-crsng.gc.ca) to Louis Bernatchez, a grant from the Mohamed bin Zayed (MBZ) Species Conservation Fund (Project 172514955; https://www.speciesconservation.org) and grant number 688MIGRD94 to Iraj Hashemzadeh Segherloo by Shahr-e-Kord University (www.sku.ac.ir), and University of Basrah. Sampling procedures comply with the current laws of Islamic Republic of Iran.

## Data availability

all haplotypes produced in this study have been deposited to the GenBank under accession nos ON203035-ON203041.

